# MiXcan: a Framework for Cell-Type-Specific Transcriptome-Wide Association Studies with an Application to Breast Cancer

**DOI:** 10.1101/2022.03.15.484509

**Authors:** Xiaoyu Song, Jiayi Ji, Joseph H. Rothstein, Stacey E. Alexeeff, Lori C. Sakoda, Adriana Sistig, Ninah Achacoso, Eric Jorgenson, Alice S. Whittemore, Robert J. Klein, Laurel A. Habel, Pei Wang, Weiva Sieh

## Abstract

Human bulk tissue samples comprise multiple cell types with diverse roles in disease etiology. Conventional transcriptome-wide association study (TWAS) approaches predict gene expression at the tissue level from genotype data, without considering cell-type heterogeneity, and test associations of the predicted tissue-level gene expression with disease. Here we develop MiXcan, a new TWAS approach that predicts cell-type-specific gene expression levels, identifies disease-associated genes via combination of cell-type-specific association signals for multiple cell types, and provides insight into the disease-critical cell type. We conducted the first cell-type-specific TWAS of breast cancer in 58,648 women and identified 12 transcriptome-wide significant genes using MiXcan compared with only eight genes using conventional approaches. Importantly, MiXcan identified genes with distinct associations in mammary epithelial versus stromal cells, including three new breast cancer susceptibility genes. These findings demonstrate that cell-type-specific TWAS can reveal new insights into the genetic and cellular etiology of breast cancer and other diseases.

## Introduction

Transcriptome-wide association studies (TWAS) aim to identify genes that are associated with disease through their genetically regulated expression levels.[1, 2] Conventional TWAS approaches such as PrediX-can [1] predict tissue-level gene expression from genetic variants using models trained on transcriptomic and genomic data from bulk tissue samples, and test associations between the predicted tissue-level expression and disease. By reducing the multiple testing burden from millions of variants to thousands of genes, TWAS can improve the power of genome-wide association studies (GWAS) while providing biological insights into the genes and regulatory mechanisms underlying disease. However, conventional TWAS approaches do not account for cell-type heterogeneity in the composition of bulk tissue samples or the effects of expression quantitative trait loci (eQTL), which can reduce the accuracy of expression prediction models and obscure disease associations, particularly when the most mechanistically relevant cell type for the disease is a minor cell type in the tissue.[3]

Breast carcinoma is a common and highly heritable cancer that arises from epithelial cells, which line the ducts and lobules that produce milk during lactation.[4, 5] Human mammary tissue has highly variable cell composition. Visualized on mammography, breast composition can range from extremely dense (light), reflecting a high proportion of fibroglandular tissue, to almost entirely fatty (dark), reflecting a high proportion of adipose tissue.[6, 7] Whereas higher mammographic density is associated with increased risk of breast cancer, a higher amount of nondense fatty tissue is associated with decreased risk, indicating disparate roles of the different cellular components of mammary tissue in carcinogenesis.[8, 9, 10] Breast cancer susceptibility loci identified by prior GWAS[11, 12, 13] and TWAS[14, 15, 16] approaches that do not account for cell-type heterogeneity explain only a fraction of the familial relative risk. Disentangling the distinct effects of gene expression in mammary epithelial cells from other cell types through cell-type-specific analysis could lead to new gene discoveries and biological insights.

To our knowledge, no statistical methods currently exist for conducting cell-type-specific TWAS using GWAS data. Single-cell sorting and transcriptome profiling are costly, and large reference panels with both single-cell transcriptomic and genomic data are not yet widely available for training robust expression prediction models from genotype data. Recent studies of bulk tissue transcriptomic data have used computational estimates of cell composition to evaluate cell-type-specific effects. The Genotype-Tissue Expression (GTEx [17]) consortium estimated cell type enrichment scores in bulk tissue samples using the xCell[18] method and tested for interactions between genotype and xCell scores in linear regression models of gene expression to identify interaction eQTLs (ieQTLs).[19] The breast was among the human tissues with the most ieQTLs, specifically involving mammary epithelial cells and adipocytes,[19] highlighting the potential for new methods that harness cell-type-specific genetic regulation of expression to improve the power of breast cancer TWAS. Methods that integrate bulk tissue data with single-cell reference profiles to estimate cell-type-specific gene expression have also been proposed to study cell-type-specific disease associations.[20, 21] However, these methods all require transcriptomic data from the disease-relevant tissue and cannot be applied to existing GWAS datasets to perform TWAS in large populations.

Here we present MiXcan, a new statistical framework for conducting cell-type-specific TWAS using GWAS data. MiXcan builds cell-type-specific gene expression prediction models through decomposition of bulk tissue data, identifies disease-associated genes via combination of signals from cell-type-specific association analyses of multiple cell types, and provides insight into the cell type responsible for the disease association. We show that MiXcan improves the prediction accuracy of gene expression at the tissue level compared with conventional approaches in an independent validation dataset. Simulation studies show that MiXcan controls the type I error, and provides higher power than conventional TWAS approaches when disease associations are driven by a minor cell type (e.g. mammary epithelial cells) rather than the predominant cell type in a tissue, or have opposite directions in different cell types. We apply MiXcan to conduct the first cell-type-specific TWAS of breast cancer risk in 31,716 cases and 26,932 controls, and report three new susceptibility genes (ZNF703, TMEM245, and PSG4) with evidence of distinct associations in mammary epithelial versus stromal cells that were not detected by prior TWAS nor GWAS. These findings provide a proof a concept that cell-type-specific TWAS can reveal new insights into the genetic and cellular etiology of breast cancer and other diseases.

## Results

### MiXcan framework

We developed the MiXcan framework for conducting cell-type-specific TWAS **(Fig. 1)**. First, the proportion of the key cell type in bulk tissue samples is estimated using the xCell[18] score as a prior (see **Methods**). Second, the bulk tissue gene expression is decomposed into its cell-type-specific components and joint penalized regression is used to model the association of genetic variants (SNPs) with gene expression (exp) for each cell type. Third, the regression coefficients (SNP weights) are compared to determine whether the prediction models are cell-type-specific or nonspecific. Association tests of the composite null hypothesis that there is no association between the genetically regulated gene expression level in any cell type with the disease versus the alternative hypothesis that there is an association in at least one cell type, are then performed using the cell-type-specific models when available and combining the resulting p-values across different cell types with the Cauchy[22] method, or using the same prediction model for all cell types similar to PrediXcan. Genes that are associated with disease are identified using the MiXcan combined p-values and applying an appropriate significance threshold to control the family-wise error rate (FWER) or false discovery rate (FDR) for the number of genes tested. For significant genes, comparison of the cell-type-specific MiXcan results provides further insight into the cell type(s) likely to be responsible for the disease association.

**Figure 1:**
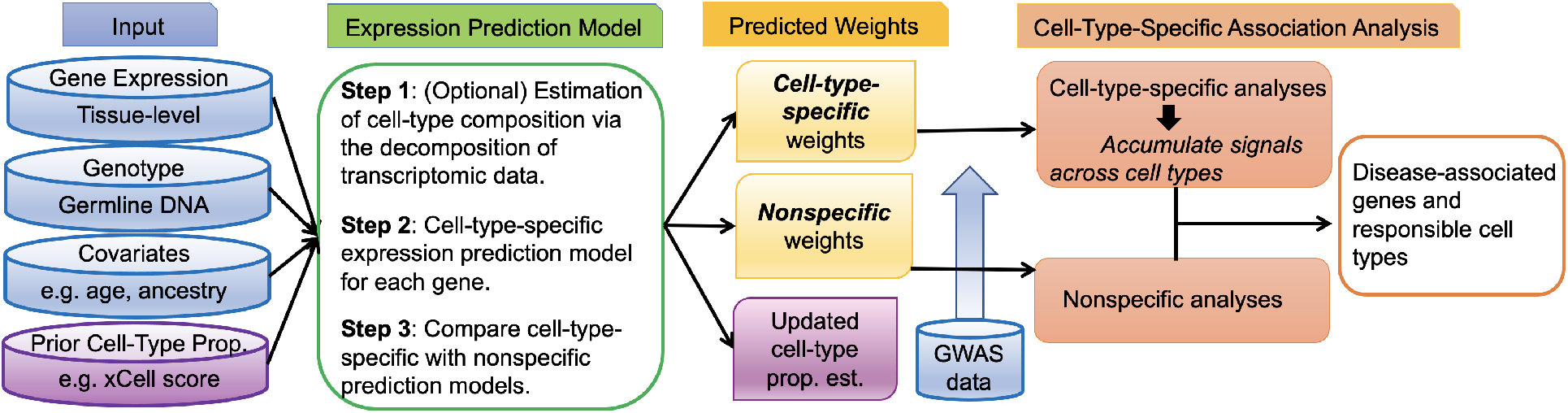
MiXcan framework. MiXcan estimates cell-type composition using transcriptomic data, builds cell-type-specific and nonspecific prediction models of gene expression, identifies disease-associated genes by aggregating the signals across different cell types, and provides insight into the cell type responsible for the disease association.

While the MiXcan framework is general, its performance depends on the cell types under consideration and the available training data. As the number of cell types increases, the number of parameters increases and the accuracy of the model decreases. At present, given the limited sample size of transcriptomic and genomic datasets available for most human tissues through public repositories such as GTEx, it is practical to consider only two categories of cells using MiXcan and to focus on the most relevant cell type for the disease versus the other cell types. To identify genes for breast cancer, we developed MiXcan epithelial and stromal (nonepithelial) cell models trained using bulk mammary tissue transcriptomic and genomic data available for 125 European ancestry (EA) women in GTEx v8.

### Prediction performance

MiXcan estimates of the epithelial cell proportion were highly correlated with the xCell[18] epithelial cell enrichment score, with Pearson correlations of 0.90 and 0.89 in normal mammary tissue samples from EA women in GTEx (N=125) and TCGA (N=103), respectively **(Supplementary Fig. 1)**. Overall, MiXcan epithelial cell proportion estimates were more highly correlated with the expression levels of 126 genes in the xCell epithelial cell gene signature (median r of 0.54 in GTEx and 0.60 in TCGA samples) than were the xCell scores themselves (median r of 0.36 in GTEx and 0.39 in TCGA samples) indicating that the MiXcan bagging approach can improve cell proportion estimation **(Supplementary Fig. 1)**.

The accuracy of MiXcan and PrediXcan gene expression prediction models trained using GTEx data for 125 mammary tissue samples from EA women was evaluated in an independent dataset of 103 tumor-adjacent normal mammary tissue samples from EA women in TCGA **(Fig. 2)**. MiXcan estimated cell-type-specific prediction models for 5473 (84.7%) and nonspecific prediction models for 988 (15.3%) of 6461 genes that had mammary tissue-level prediction models available in PredictDB.[23] Gene expression at the tissue level was computed using MiXcan estimates of the cell-type proportion and predicted component gene expression levels. The median correlation of predicted and measured mammary tissue expression levels for the 6461 genes in the TCGA validation set was significantly higher for MiXcan compared with PrediXcan (median r of 0.41 vs. 0.10; *p*-value<2.2×10-16) models trained using the same dataset of 125 GTEx EA women. The prediction accuracy for the 5473 genes with cell-type-specific models in MiXcan was significantly better than PrediXcan (median r of 0.43 vs. 0.12; *p*<2.2×10 16), whereas the prediction accuracy for the remaining 988 genes with nonspecific models in MiXcan was the same as PrediXcan (median r of 0.08 vs. 0.08; *p*-value=1). These results indicate that allowing for cell-type-specific gene expression models increases the prediction accuracy for genes with evidence of cell-specific expression, and does not decrease the prediction accuracy for other genes compared with standard approaches for predicting tissue-level gene expression.

**Figure 2:**
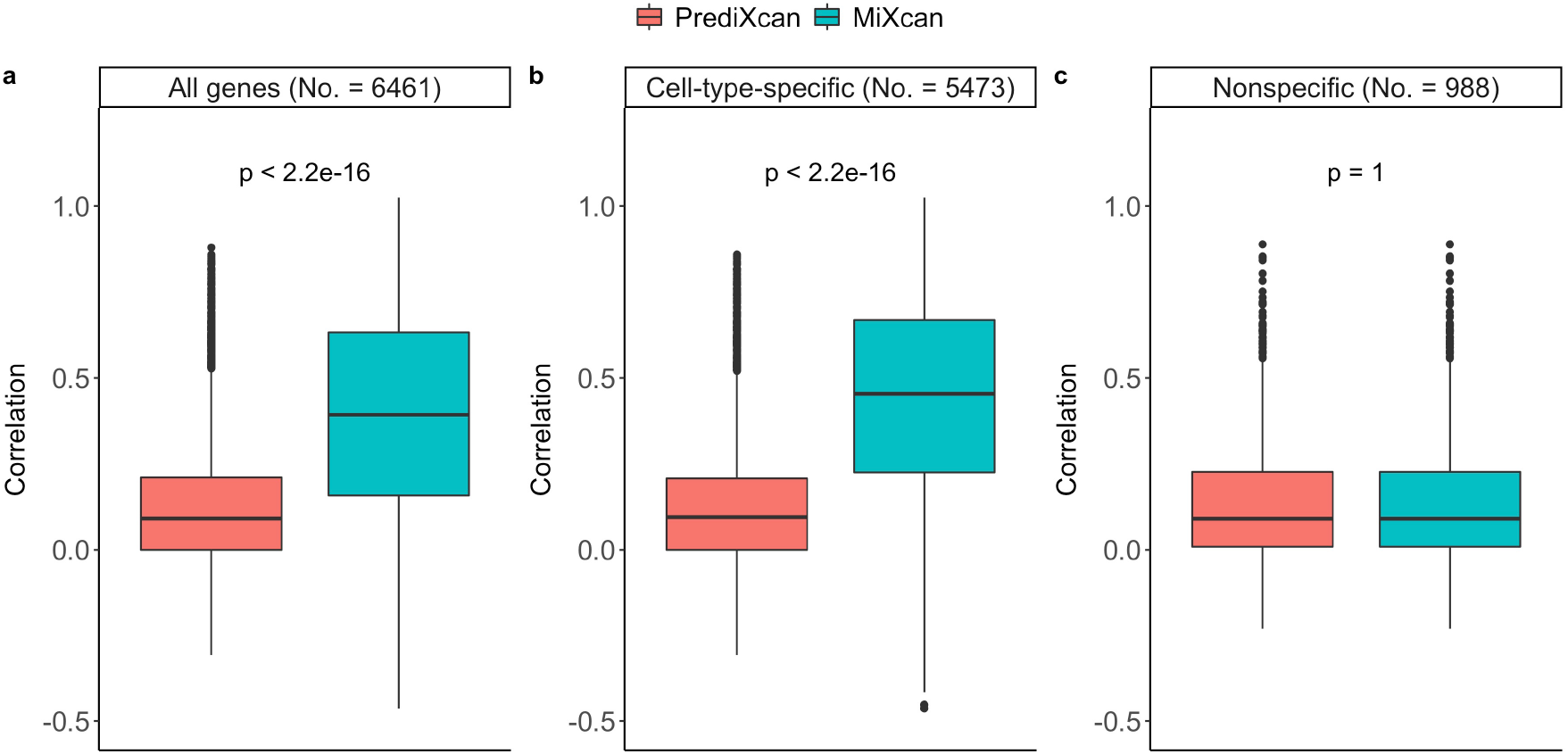
Gene expression prediction accuracy in independent validation dataset. Correlation of tissue-level gene expression measurements with predictions using MiXcan or PrediXcan in an independent dataset of adjacent normal mammary tissue samples from 103 European ancestry women with breast cancer in TCGA. MiXcan and PrediXcan prediction models were developed using mammary tissue samples from 125 European ancestry women in GTEx v8 for 6461 genes (**a**), including 5473 genes with cell-type-specific (**b**) and 988 genes with nonspecific (**c**) MiXcan prediction models.

To examine potential sources of the gain in prediction accuracy, three additional approaches were compared with MiXcan and PrediXcan **(Supplementary Fig. 2)**. The median correlation of predicted and measured mammary tissue expression levels for all 6461 genes in the TCGA validation set was slightly higher for PredictDB (r=0.12) elastic-net models trained using 337 GTEx EA men and women compared with PrediXcan (r=0.10) trained using 125 EA women indicating modest gains from the inclusion of 212 EA men in the training dataset. Accounting for cell composition using penalized regression models including interactions of SNPs with the xCell epithelial cell score (xCell Interaction; r=0.20) or MiXcan cell proportion (MiXcan_0_; r=0.38) led to substantial gains in prediction accuracy. Symmetric estimation of cell-type-specific prediction models employed in MiXcan (r=0.41) further improved performance. Importantly, whereas standard interaction models require estimates of cell-type composition, which often are unavailable for the tissue of interest in GWAS of human diseases, MiXcan prediction models can be applied directly to GWAS genotype data to perform cell-type-specific TWAS.

### Simulation studies

To evaluate type I error and power of MiXcan association tests, datasets were simulated (see **Methods**) under a broad range of realistic models for the associations of genetic variants with gene expression (SNP-exp) and gene expression with disease (exp-disease). MiXcan predicted gene expression levels with higher accuracy than PrediXcan in the presence of cell-type heterogeneity of SNP-exp associations, while maintaining comparable accuracy in the absence of cell-specific effects **(Supplementary Fig. 3)**, consistent with results in the independent TCGA validation dataset **(Fig. 2)**.

Association test results for MiXcan using either the true or misspecified cell-type proportions were compared with PrediXcan in simulated datasets **(Fig. 3)**. The type I error was well controlled for MiXcan and PrediXcan under all simulated data scenarios **(Fig. 3 col. 1)**. When SNP-exp associations were homogeneous in the two cell types **(Fig. 3a)**, the power was similar for MiXcan and PrediXcan whether the exp-disease associations were homogenous or heterogeneous across cell types. Differences in the mean gene expression level between the two cell types that were not determined by SNP-exp associations **(Fig. 3a-b)** did not impact the power of MiXcan and PrediXcan indicating robustness to differential expression that is not regulated by genetic variants. MiXcan results using the true and misspecified values of the cell-type proportion were comparable indicating robustness to noisy estimates of the cell-type proportion **(Fig. 3; Supplementary Fig. 3)**.

**Figure 3:**
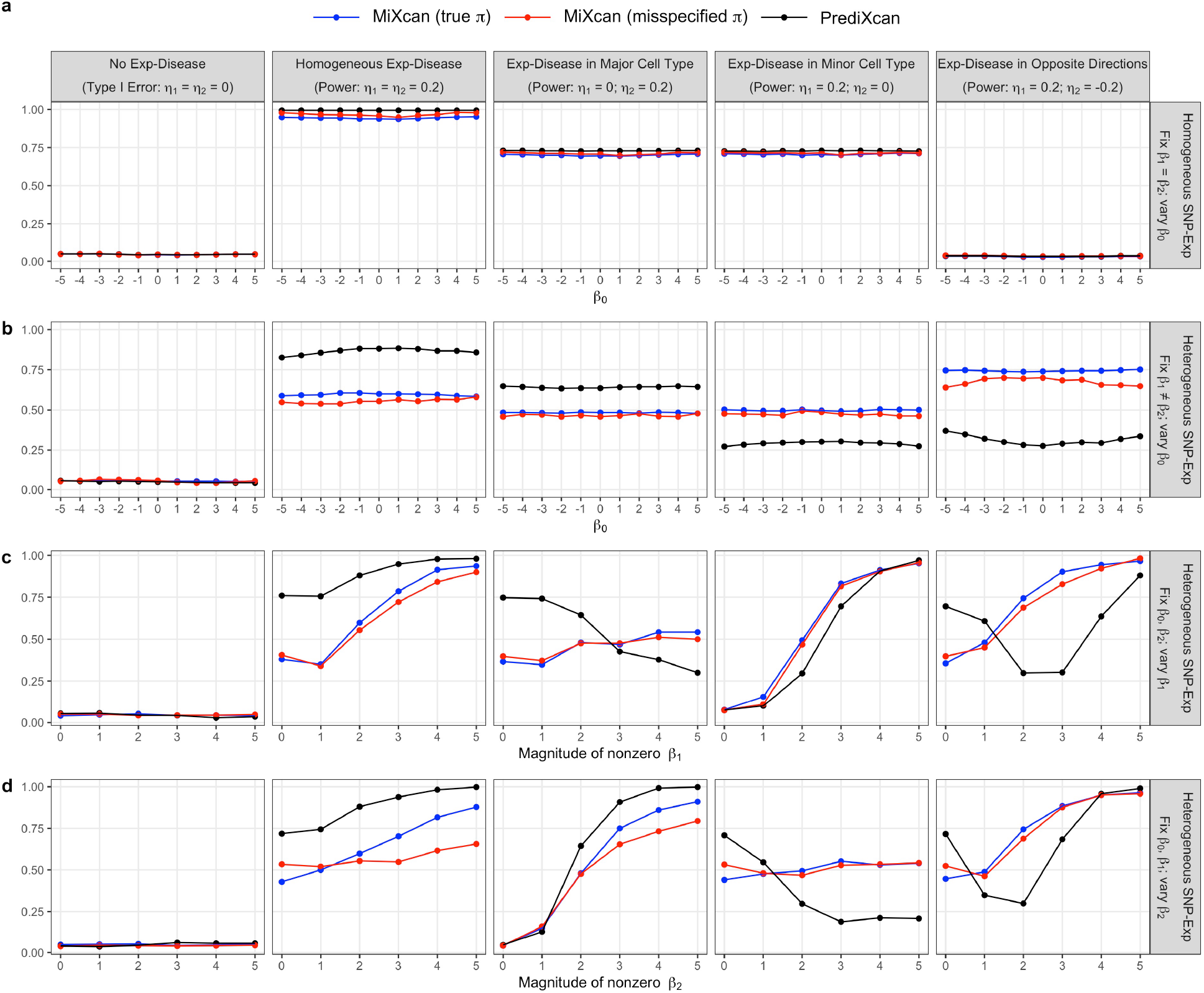
Simulation studies. Type I error and power of MiXcan and PrediXcan to detect associations of tissue-level gene expression with the disease in independent test sets (N=500) simulated under a range of realistic data scenarios. MiXcan results also are shown when the minor cell-type proportion is misspecified, with a correlation of 0.9 and mean shift of 5% from the true value. Gene expression levels were modeled by *u* = *β*_0_ + ***β***_1_*x* + *e*_*u*_ in the minor cell type, *v* = ***β***_2_*x* + *e*_*v*_ in the major cell type, and *y* = *πu* + (1 − *π*)*v* at the tissue level, where *π* denotes the minor cell-type proportion, *β*_0_ denotes the mean difference of the gene expression levels in the two cell types, and ***β***_1_ and ***β***_2_ denote the weights for the association of SNPs X with gene expression levels in the minor and major cell types, respectively. The disease D was modeled by logit *P*(*D* = 1) = *η*_0_ + *η*_1_*u* + *η*_2_*v* where *η*_1_ and *η*_2_ denote the associations of the gene expression levels with disease in the two cell types, respectively. (**a**) Homogeneous SNP-exp associations (***β***_1_ = ***β***_2_) in the two cell types, varying the mean difference in gene expression levels between the two cell types (*β*_0_). Heterogeneous SNP-exp associations (***β***_1_ ≠ ***β***_2_) in the two cell types, varying the: (**b**) mean difference in gene expression levels between the two cell types (*β*_0_); (**c**) magnitude of the SNP-exp association in the minor cell type (***β***_1_); and (**d**) magnitude of the SNP-exp association in the major cell type (***β***_2_).

When SNP-exp associations were heterogeneous in the two cell types **(Fig. 3b-d)**, the relative power of MiXcan and PrediXcan depended on the mechanisms of the exp-disease and SNP-exp associations. PrediX-can was generally more powerful than MiXcan when the exp-disease association was either homogeneous across cell types **(Fig. 3 col. 2)** or present only in the major cell type **(Fig. 3 col. 3)**. However, MiXcan was generally more powerful than PrediXcan when the exp-disease association was present only in the minor cell type **(Fig. 3 col. 4)** or had opposite directions in the two cell types **(Fig. 3 col. 5)**.

As the strength of the SNP-exp association increased in the same cell type as the exp-disease association, the power increased for both PrediXcan and MiXcan **(Fig. 3c col. 4; Fig. 3d col. 3)**. However, as the strength of the SNP-exp association increased in a different cell type from the exp-disease association, the power decreased for PrediXcan but not MiXcan **(Fig. 3c col. 3; Fig. 3d col. 4)**. When the exp-disease association had opposite directions in the two cell types, the power was U-shaped for PrediXcan but increased for MiXcan as the strength of the SNP-exp association increased in either cell type **(Fig. 3c-d col. 5)**. These patterns show that different association signals in the two cell types can cancel each other out in PrediXcan, which averages their effects, but are aggregated across cell types in MiXcan thereby preserving power to detect associations due to the minor cell type or that differ across cell types.

In addition to providing valid tissue-level tests of the association of predicted gene expression with disease by aggregating the cell-type-specific signals, MiXcan provides information for each cell type separately. Simulation studies showed that the type I error was well controlled for MiXcan cell-type-specific tests when no exp-disease association was present **(Supplementary Fig. 4 col. 1)**. When SNP-exp associations were homogeneous across cell types **(Supplementary Fig. 4a)**, the expression prediction model and association test results were the same for all cell types. Thus, cell-type-specific inferences can only be made in the heterogeneous SNP-exp setting **(Supplementary Fig. 4b-d)**, when cell-type-specific expression prediction models are available in MiXcan. When the exp-disease association was present in both cell types in the same or opposite directions **(Supplementary Fig. 4 cols. 2 & 5)**, the power of the cell-type-specific tests was similar when the SNP-exp associations had similar magnitude (regardless of direction) and increased as the magnitude of the SNP-exp association increased. When the exp-disease association was present in only one cell type **(Supplementary Fig. 4 cols. 3-4)**, the power was always highest in this cell type, but the association signal was shared to some degree with the uninvolved cell type. This correlation of the cell-type-specific results arises from the joint estimation of the SNP weights for the cell-type-specific expression prediction models in MiXcan. Therefore, we recommend using the combined p-value for all cell types to make inferences regarding whether gene expression is significantly associated with disease in any cell type in the tissue, and the cell-type-specific results to compare the evidence that different cell types are involved for significant genes.

Finally, we evaluated the impact of the sample size of the training dataset on the type I error and power of MiXcan association tests in simulation studies **(Supplementary Fig. 5)**. As the training dataset increased from 100 to 300 samples, the power of association studies with 500 samples increased while the type I error remained well controlled. Prediction models trained using only 100-150 samples provided reasonable power for gene identification.

### Cell-type-specific TWAS of breast cancer

We applied MiXcan to conduct the first cell-type-specific TWAS of breast cancer in 58,648 EA women (31,716 cases and 26,932 controls) who were genotyped using the OncoArray[24] in the Discovery, Biology, and Risk of Inherited Variants in Breast Cancer (DRIVE) GWAS available in dbGaP (accession number phs001265.v1.p1). Transcriptome-wide significance was determined using the Bonferroni-corrected threshold of 7.7×10-6 to account for the 6461 genes tested. MiXcan identified 12 genes whose predicted expression levels in mammary tissue were significantly associated with breast cancer risk **(Table 1, Fig. 4)**, and 82 suggestive genes at the false discovery rate (FDR) of 0.10 using the Benjamini-Hochberg procedure **(Supplementary Table 1)**. In comparison, PrediXcan (trained using the same 125 EA female mammary tissue samples as MiXcan) identified only 8 significant genes (*p*-value<7.7×10-6), and 31 suggestive genes (FDR<0.10). The genomic inflation factor, *λ*_1000_,[25, 26] was 1.008 for MiXcan and 1.011 for PrediXcan indicating that the type I error was well controlled.

**Table 1.**
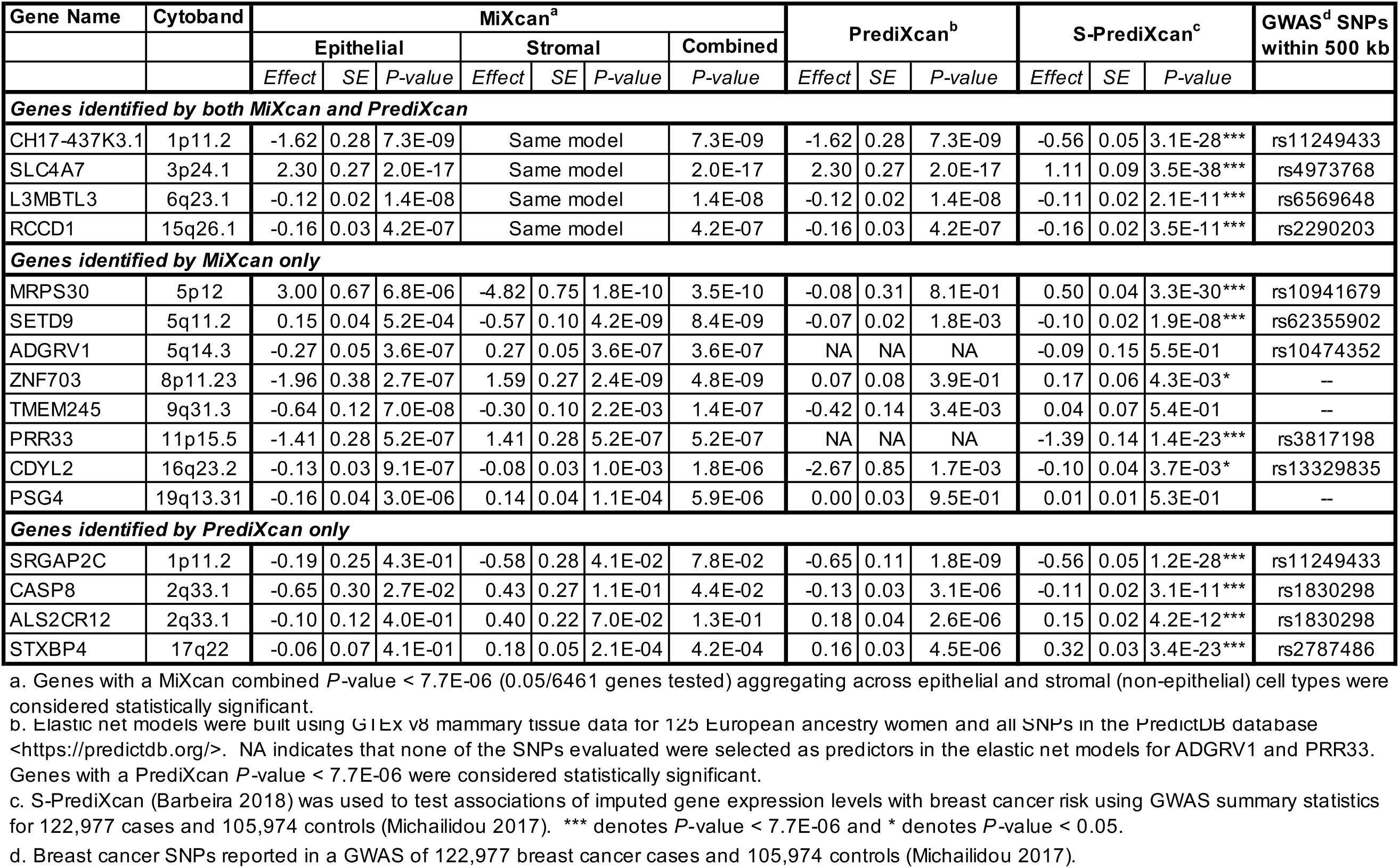
Genes significantly associated with breast cancer risk using the MiXcan or PrediXcan approaches in 58,648 women, and the corresponding S-PrediXcan and GWAS results in a larger sample of 228,951 women of European ancestry.

**Figure 4:**
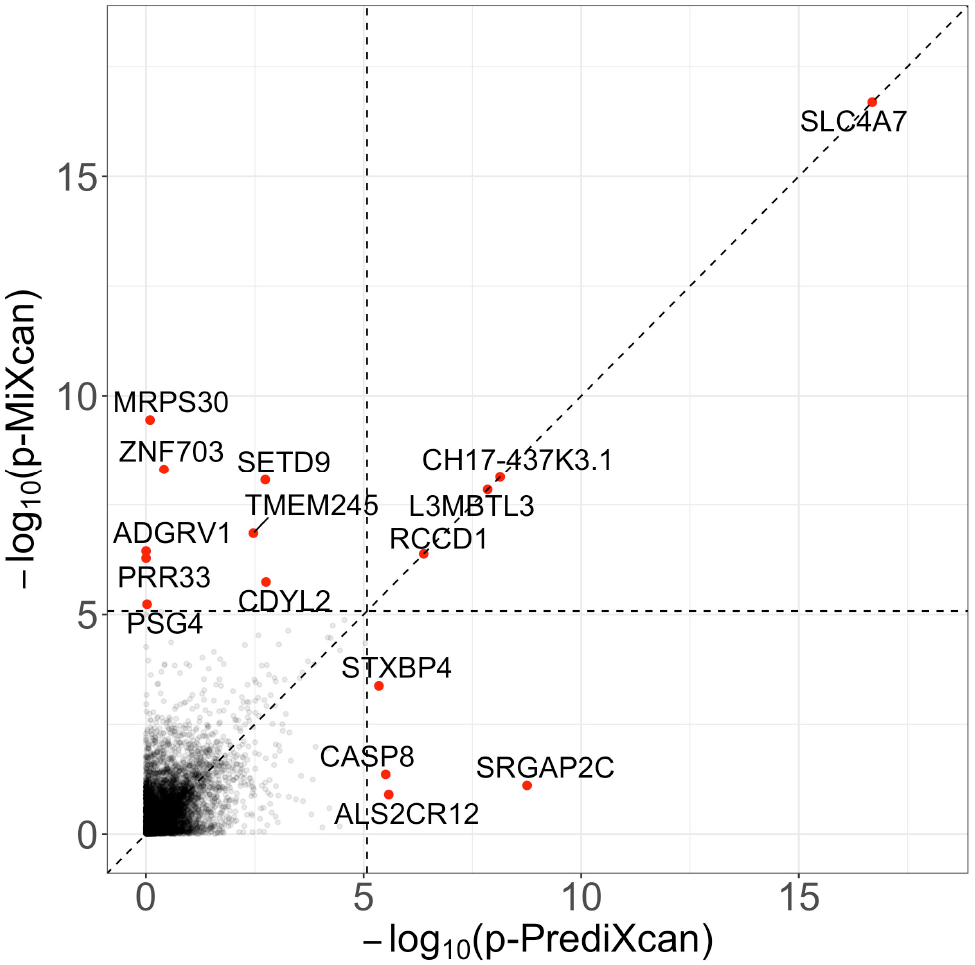
Transcriptome-wide association studies. Genes significantly associated with breast cancer using MiXcan or PrediXcan at *p*-value < 7.7 × 10-6 applying a Bonferroni correction for the 6461 genes tested in 31,716 breast cancer cases and 26,932 controls of European ancestry from the DRIVE study (dbGaP phs001265.v1.p1).

Four significant genes (CH17-437K3.1, SLC4A7, L3MBTL3, and RCCD1) were identified by both MiXcan and PrediXcan **(Table 1)**. MiXcan did not estimate cell-type-specific prediction models for these genes, and a homogeneous prediction model was used yielding the same results as PrediXcan. All four genes were near (<500 kb) breast cancer SNPs previously identified by GWAS.[11] Follow-up analyses in a larger sample of 228,951 EA women (122,977 cases and 105,974 controls) in the Breast Cancer Association Consortium (BCAC) and DRIVE studies using GWAS summary statistics[11] and S-PrediXcan[27] models for mammary tissue confirmed that the tissue-level expression for all four genes were significantly (*p*-value <7.7×10-6) associated with breast cancer risk.

Eight genes were identified by MiXcan but not PrediXcan **(Table 1)**. Six of these genes (MRPS30, SETD9, ADGRV1, ZNF703, PRR33, and PSG4) showed different directions of association with breast cancer in epithelial vs. stromal (nonepithelial) cells. In MiXcan the signals from the two cell types were aggregated, whereas in PrediXcan they canceled each other out reducing the tissue-level signal. Notably, for ADGRV1 and PRR33, no SNPs in the training dataset were predictive of gene expression at the tissue level because of the mixture of the different cell types, and the PrediXcan analysis could not be performed. Two genes (TMEM245 and CDYL2) showed the same directions of association, with stronger effects in epithelial vs. stromal cells. These results indicate that MiXcan may be more powerful than PrediXcan in the presence of cell-type heterogeneity of genetically regulated expression and when the disease association is present in a minor cell type, e.g. mammary epithelial cells, rather than the predominant cell type in the tissue.

Importantly, MiXcan uniquely identified three novel breast cancer susceptibility genes (ZNF703, TMEM245, and PSG4) that were not previously implicated by breast cancer GWAS[11, 12, 13] nor TWAS[14, 15, 16] **(Table 1)**. For ZNF703, gene expression was associated with increased breast cancer risk in stromal cells (*p*-value=2.4×10-9) and decreased risk in epithelial cells (*p*-value=2.7×10-7). Because adipocytes are the predominant stromal cell type in mammary tissue, we also performed follow-up analyses using the S-PrediXcan subcutaneous fat model and discovered a highly significant (*p*-value=2.1×10-20) association of ZNF703 with increased breast cancer risk in BCAC/DRIVE, consistent with the MiXcan stromal cell results. For TMEM245 and PSG4 the signal was stronger in epithelial cells, which are a minor cell type in mammary tissue and may explain why their tissue-level expression was not significantly associated with breast cancer risk. MiXcan also identified two breast cancer genes (ADGRV1 and CDYL2) at previously reported GWAS loci[11] that had different associations in epithelial and stromal cells and were not detected in prior TWAS.

Four genes (SRGAP2C, CASP8, ALS2CR12, and STXBP4) were identified by PrediXcan but not MiXcan **(Table 1)**. There was high correlation between the predicted tissue-level expression of CH17-437K3.1 (also identified by MiXcan) and SRGAP2C at 1p11.2 (r=0.95) and CASP8 and ALS2CR12 at 2q33.1 (r=-0.97) indicating that these associations may represent only two independent loci. All four genes identified by PrediXcan only were located near breast cancer SNPs previously identified by GWAS,[11] and associations of the mammary tissue-level expression with breast cancer risk were also found by S-PrediXcan analyses of the BCAC/DRIVE data. MiXcan estimated cell-type-specific prediction models for these four genes, and different associations with breast cancer risk in epithelial and stromal cells that did not reach statistical significance in part because of the larger number of model parameters compared with PrediXcan. However, the cell-type-specific MiXcan results for STXBP4 suggest that stromal cells (estimated effect=0.18; *p*-value=2.1×10-4) may play a more important role than epithelial cells (estimated effect=-0.06; *p*-value=0.41) in driving the positive association of tissue-level expression with breast cancer risk.

Finally, we compared the TWAS results using MiXcan and PrediXcan with publicly available PredictDB[23] elastic-net models trained using GTEx mammary tissue data for 212 EA men in addition to 125 EA women **(Supplementary Fig. 6)**. There was substantial overlap of the genes detected by PrediXcan and PredictDB as expected, although PredictDB detected a larger number of genes perhaps because the larger training dataset enabled more accurate prediction models for genes that have similar expression patterns in male and female mammary tissue. Cell-type-specific TWAS using MiXcan identified five genes (ADGRV1, ZNF703, TMEM245, CDYL2, and PSG4), including three novel breast cancer susceptibility genes, that were not identified by either PrediXcan or PredictDB.

## Discussion

MiXcan is a new statistical framework for conducting cell-type-specific TWAS using GWAS data. In contrast to standard TWAS methods, MiXcan builds cell-type-specific prediction models for the genetically regulated component of gene expression and performs association tests taking into consideration the signals from multiple cell types. We have shown that MiXcan improves the prediction accuracy of gene expression at the tissue level in both simulation studies and an independent validation dataset, and improves the power to detect disease associations that are driven by a minor cell type or are heterogeneous between cell types compared with standard approaches. We applied MiXcan to perform the first cell-type-specific TWAS of breast cancer risk and discovered three new susceptibility genes (ZNF703, TMEM245, and PSG4) with evidence of distinct associations in mammary epithelial versus stromal cells that were not detected by prior GWAS[11, 12, 13] nor TWAS.[14, 15, 16] These findings provide a proof of concept that cell-type-specific TWAS can reveal novel insights into the genetic and cellular etiology of human diseases.

Several recent studies have explored methods for performing cell-type-specific association analyses when the tissue-level transcriptomic or epigenomic data are available for all subjects.[19, 28, 29, 30, 31] In this study, MiXcan enables cell-type-specific TWAS using existing GWAS datasets without transcriptomic and cell composition data from the disease-relevant tissue, which can be infeasible to collect in large populations. Specifically, to reveal biologically meaningful association signals and address potential confounding due to the cell composition of bulk tissue samples, MiXcan evaluates the composite null hypothesis that there is no association between the genetically regulated gene expression level in any cell type with the disease. By carefully modeling cell-type-specific expression, MiXcan is more powerful than PrediXcan when disease associations are driven by a minor cell type or have opposite directions in different cell types. However, when the association of gene expression with disease is similar in all cell types or driven by the major cell type, then conventional TWAS approaches using more parsimonious tissue-level expression prediction models that assume cell-type homogeneity can be more powerful. Thus, these two TWAS approaches are complementary, and additional cell-type-specific analyses are especially valuable for diseases where cell-type heterogeneity and a minor cell of origin are hypothesized, as for breast carcinoma and many other human diseases.

To construct cell-type-specific expression prediction models, MiXcan uses the xCell[18] cell-type enrichment score in the training data as prior information. While estimates from other approaches[20, 21, 32, 33, 34, 35] could also be used as priors, xCell is among the most widely used. Building upon the priors, MiXcan fits mixture models for the expression levels of the epithelial cell signature genes in the training data to improve estimation of the cell-type proportion. By incorporating better estimates of the cell-type proportion, and penalizing all cell types equally, MiXcan improves the accuracy of the expression prediction models, as well as the power and type I error of the downstream association tests. Compared with standard interaction models that include interaction effects between cell-type enrichment scores and genetic variants,[19] an important advantage of MiXcan is that the predicted values correspond to the cell-type-specific genetically regulated gene expression levels, which are directly interpretable and biologically meaningful. Moreover, standard interaction models require cell composition information, which is often unavailable for the tissue of interest, whereas MiXcan prediction models can be applied directly to GWAS data without transcriptomic or cell composition data to interrogate genetic associations within the disease-relevant tissue and cell type context.

Although the MiXcan framework is generalizable, the model building procedure requires knowledge of the disease-critical cell types and tissue. The present prediction models were developed specifically for human mammary tissue with a focus on distinguishing epithelial and stromal (nonepithelial) cells, which have distinct roles in breast carcinogenesis.[8, 9, 10] The MiXcan framework can be extended naturally to model more than two cell types, but a balance between the model complexity and the sample size should be considered. As larger training sets with bulk tissue transcriptomic and genomic data become available, a careful evaluation of its analytical performance can be performed for less common cell types. Currently, human single-cell transcriptome profiling efforts[36, 37] are underway, although the available sample sizes are still too small for direct training of robust prediction models. Future studies can investigate the integration of single-cell transcriptome profiles to improve estimation of cell composition in bulk tissue samples[35, 34] or to provide an initial estimate of SNP weights, which could be used to tune separate penalty terms for different SNPs in adaptive elastic-net models.[38]

Human mammary tissue has variable cell composition and numerous eQTLs with distinct effects in epithelial cells and adipocytes, which are a major stromal cell type in the breast.[19] Cell-type-specific TWAS using MiXcan mammary tissue models applied to publicly available GWAS data identified three new breast cancer susceptibility genes that were associated with disease risk through their genetically regulated expression levels in normal mammary epithelial or stromal cells. ZNF703 (zinc finger protein 703) is an oncogene that is commonly amplified in luminal B breast tumors, and has been shown to regulate genes involved in proliferation, invasion, and an altered balance of progenitor stem cells.[39, 40, 41] To our knowledge, common germline variants in ZNF703 have not previously been implicated in breast cancer risk. Our finding that genetic upregulation of ZNF703 in normal mammary stromal cells (predominantly adipocytes) was associated with increased breast cancer risk in 58,648 women was confirmed by a highly significant (*p*-value=2.1×10 20) association of tissue-level expression predicted using S-PrediXcan subcutaneous fat models in 228,951 EA women, which has not previously been reported to our knowledge. Notably, the mammary tissue-level results for ZNF703 did not reach transcriptome-wide significance, underscoring the importance of accounting for cell-type heterogeneity to elucidate disease etiology. TMEM245 (transmembrane protein 245) is the host gene for microRNA 32, which has been shown to promote proliferation and suppress apoptosis of breast cancer cells.[42] Relatively little is known about PSG4 (pregnancy specific beta-1-glycoprotein 4), a member of the carcinoembryonic antigen gene family that may play a role in regulation of the innate immune system.[43]

In conclusion, the MiXcan framework enables cell-type-specific TWAS using prediction models for genetically regulated gene expression that allow for differences across cell types in the disease-relevant tissue. *MiXcan* mammary tissue models are available at https://github.com/songxiaoyu/MiXcan and can be applied to GWAS genotype data to identify genes associated with complex traits through their expression levels in epithelial or stromal cells. *MiXcan* software is also freely available to facilitate training prediction models for additional tissues and cell types, and conducting cell-type-specific TWAS. MiXcan prediction models had excellent performance in an independent validation dataset, and identified new breast cancer susceptibility genes in the first cell-type-specific TWAS of breast cancer. These findings provide a proof of concept that cell-type-specific TWAS are feasible using existing bulk tissue training datasets and GWAS data, and can lead to the discovery of new disease genes and cellular mechanisms. Future research is needed to develop MiXcan models for other human tissues and cell types, and to extend the MiXcan framework for application to GWAS summary statistics to enable the broad application of cell-type-specific TWAS to improve our understanding of the genetic and cellular mechanisms underlying human diseases.

## Methods

### MiXcan framework

We first summarize PrediXcan,[1] an established tissue-level TWAS framework, and then present the MiXcan cell-type-specific TWAS framework. Let *y*_*i*_ denote the measured expression level of a gene in the bulk tissue sample *i* ∈ (1, …, *N*), ***x***_*i*_ denote the genetic variants (e.g. SNPs) used to predict gene expression, and ***z***_*i*_ denote the non-genetic covariates (e.g. age).

### PrediXcan tissue-level TWAS framework

PrediXcan uses a linear additive model to characterize the gene expression level:

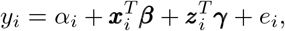

where *e*_*i*_∼*N*(0, *σ*^2^) is the model error, 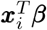 is the genetically determined component of gene expression, and 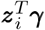 is the non-genetically determined component. This model can be estimated with the elastic-net method,[44] which maximizes the following penalized log-likelihood function:

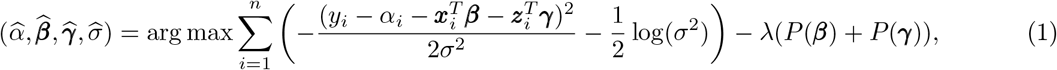

where 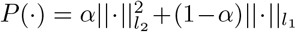 is the elastic-net penalty function with the mixing parameter *α* ∈ (0, 1). In PrediXcan, *α* is set at 0.5 and λ is selected via 10-fold cross validation.(CV) [45] The estimated SNP weights 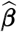, can be used to predict the genetically regulated gene expression levels by 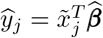 where 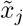 denotes the SNP genotypes of the GWAS subject *j* ∈ (1, …, *M*). Then, the association of *ŷ*_*j*_ with the phenotype (e.g. disease status) *d*_*j*_ can be evaluated using a generalized linear model *g*(*d*_*j*_) = *η*_0_ + *ŷ*_*j*_*η*_1_, where *g*(.) is a link function. The null and alternative hypotheses, *H*_0_: *η*_1_ = 0 vs. *H*_*A*_: *η*_1_ ≠ 0, test whether the genetically regulated gene expression at the tissue level is associated with the phenotype.

### MiXcan cell-type-specific expression prediction model

MiXcan extends upon PrediXcan to enable cell-type-specific TWAS. In this section, we build the prediction models for the cell-type-specific genetically regulated gene expression levels. In the next section, we develop strategies for applying these prediction models in disease-association studies.

Human bulk tissue samples from solid tissue (e.g. mammary tissue) comprise a mixture of cells of different types. Let *π*_*i*_ and 1 − *π*_*i*_ be the proportions of the cell type of interest (e.g. epithelial cells) and all other cell types (e.g. stromal cells) in the *i*^*th*^ tissue sample, respectively. We assume the observed bulk tissue expression level of a given gene *y*_*i*_ is a linear combination of expression levels in the cell types. Introducing two latent variables *u*_*i*_ and *v*_*i*_ to denote the unobserved average gene expression levels in the epithelial and stromal cells, respectively, we have:

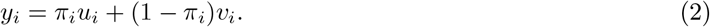

We model both *u*_*i*_ and *v*_*i*_ using the linear additive models:

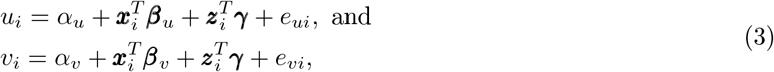

where 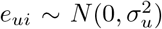 and 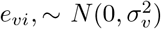 are the model errors. In Equation(3), the intercepts *α*_*u*_ and *α*_*v*_ and genetic parameters ***β***_*u*_ and ***β***_*v*_ differ between cell types, allowing for different mean expression levels and cell-type-specific effects of genetic variants on gene expression. A shared parameter ***γ*** is used for the non-genetic component ***z***_*i*_ to simplify the model as the non-genetic variables are not necessarily used in downstream analyses.

The proportion of epithelial cells ***π*** = {*π*_1_, …, *π*_*N*_} is a feature of the tissue samples, which can be jointly estimated using multiple genes, whereas 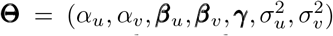 are features of each gene under investigation. Therefore, we present a step-wise procedure to first estimate ***π*** using multiple genes, and then estimate the cell-type-specific effects **Θ** for each gene.

#### Estimation of *π*

The notation in the sections above focuses on individual genes, and here we introduce additional notation to describe the joint modeling of multiple genes. Let 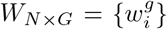 be the observed expression of *G* epithelial cell signature genes in *N* tissue samples; and 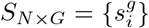 and 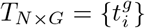 be the unobserved gene expression levels in epithelial and stromal cells, respectively. Similarly as in Equation (2), we have:

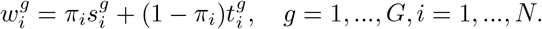

Leveraging primarily the mean differences of signature genes in epithelial and stromal cells for estimating ***π***, we model the marginal distributions of individual genes and omit the complex gene-gene correlations for computationally efficient estimation, as supported by our previous work.[46] Specifically, we assume:

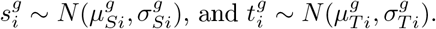

Across all *G* genes, the parameters include 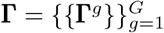, where 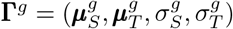.

In parallel, we also take advantage of a prior cell-type proportion estimate *h*_*i*_ based on existing tools (i.e. xCell[18] enrichment scores). We link the prior estimates *h*_*i*_ to the true *π*_*i*_ using a *Beta* distribution such that *h*_*i*_ *∼ Beta*(*π*_*i*_ *δ*, (1 − *π*_*i*_) *δ*) for some positive parameter *δ*. We have *E*(*h*_*i*_) = *π*_*i*_ and *var*(*h*_*i*_) = *π*_*i*_(1 − *π*_*i*_)*/*(*δ*+1) such that *h*_*i*_ is an unbiased estimator of the true *π*_*i*_ with variation.

We then join these two models for parameter estimation and solve the following maximization problem:

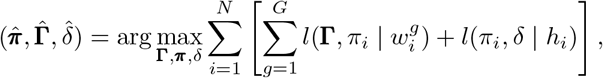

where 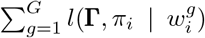 and *l*(*π*_*i*_, *δ*| *h*_*i*_) are the log-likelihood of the observed gene expression profile and cell proportion estimate of the *i*^*th*^ sample, respectively. This optimization problem is solved using an Expectation-Maximization (EM) algorithm similar to that in Petralia et al.[46]

To enhance the robustness of the estimation, we implement a bagging strategy to estimate the parameters with randomly selected bootstrap samples, and aggregate multiple estimates by calculating a tail truncated mean. This bagging strategy further stabilizes the estimates, and may also be used to investigate the consistency of ***π*** estimation.

#### Estimation of *β*_*u*_ and *β*_*v*_

Given 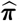, we next estimate ***β***_*u*_ and ***β***_*v*_ in Equation (3). Since *u*_*i*_ and *v*_*i*_ are unobserved, we integrate Equations (2) and (3) to have:

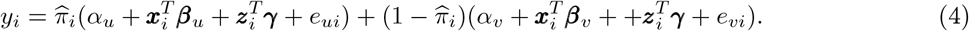

This equation can be rearranged as:

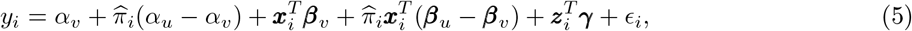

where 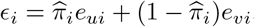. A simple strategy for estimating ***β***_*u*_ and ***β***_*v*_ is to apply elastic-net regression to Equation (5). Specifically, the elastic-net regularization is put on **(*β***_*v*_, ***β***_*u*_ − ***β***_*v*_, ***γ***)—the dependence of expression levels on genetic variants and covariates—but not on (*α*_*v*_, *α*_*u*_ − *α*_*v*_) —the mean expression levels in the two cell types. We refer to this strategy as MiXcan_0_.

One issue with MiXcan_0_ is that the two cell components in the mixture model are not treated in a symmetric manner. In other words, the penalization on ***β***_*u*_ and ***β***_*v*_ differs: ***β***_*v*_ is shrunk towards zero, while ***β***_*u*_ is shrunk towards ***β***_*v*_. This asymmetric penalization results in different models if the order of the two components is switched. To address this issue, we introduce 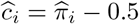 and rewrite Equation (5) as:

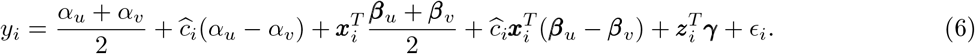

When fitting elastic-net regression to Equation (6), we include penalties on 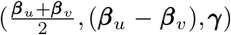, which impose the same degree of regularization on ***β***_*u*_ and ***β***_*v*_: the penalty on 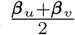 regularizes the overall sparsity of the genetic effects, while the penalty on ***β***_*u*_ − ***β***_*v*_ encourages similarities between the two components. We refer to this strategy as MiXcan.

Note, when fitting elastic-net regressions in MiXcan_0_ and MiXcan, we do not consider the varying variances of *ϵ*_*i*_ as 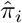 takes different values. This is because the residual variance structure has limited impact on the coefficient estimates, especially for regularized regression. In a trade-off between extensive computational costs (allowing varying residual variances) and minimal sacrifice of estimation accuracy (assuming constant variance), we chose the latter and take advantage of the fast implementation of elastic-net regression in the *glmnet* package.

#### Model Aggregation

In the prediction models, the term 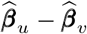 is of particular importance: a non-zero value suggests that the dependence structure between genetic variants and expression levels is cell-type-specific. Therefore, it is critical to know the selection robustness of 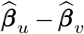. We employ a procedure similar to stability selection [47] for its evaluation. Specifically, for models that select non-zero 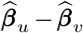, we generate *B* bootstrap samples (e.g. *B* = 200), perform ordinary least square analysis on the pre-selected variables, and record 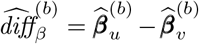 for *b* = 1, …, *B*. Only when the 95% confidence interval (CI) for 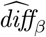 excludes 0 do we employ cell-type-specific prediction models (inferred using the complete data set). Otherwise, the same prediction model will be used for both cell types, as in Equation (1) of PrediXcan.

### Association analysis with cell-type-specific prediction models

The model building procedure in MiXcan selects cell-type-specific prediction models for some genes and nonspecific models for other genes. Nonspecific models are the same models as developed in PrediXcan, and thus can use the existing association strategy. Cell-type-specific prediction models, however, employ cell-type specific prediction weights, 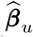 and 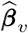, to predict expression levels from the genotype data 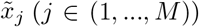, such that 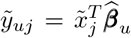 and 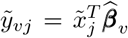. These cell-type-specific expression levels cannot be combined into tissue levels in GWAS datasets that lack cell-type proportion estimates, requiring a novel statistical framework for association analysis.

One natural idea is to infer cell-type-specific associations by directly associating the phenotype *d*_*j*_ with 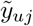 and 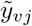, either separately, such that 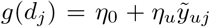 and 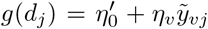, or jointly, such that 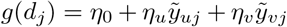. While appealing, this idea is problematic when using the predicted values. This is because the 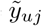 and 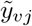 are predicted from the same genotype data with prediction weights jointly learned using the bulk tissue data. Model uncertainty is expected, and 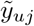 and 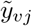 might capture leaked information from other cell types. As a result, if an association exists in one cell type, analysis in the other cell type may also capture this association, resulting in an inflated type I error for cell-type-specific association analysis. To avoid this inflation, we propose a composite hypothesis test of whether any association exists in the tissue:

> *H*_0_ : There is no association in any cell type in the tissue vs.
>
> *H*_*A*_ : There is an association in at least one cell type in the tissue.

This composite null is robust against information leakage, as the leaked values under the null are not associated with the phenotype. To perform the test, we first associate *d*_*j*_ with 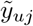 and 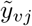, separately if the 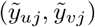 are highly correlated (e.g. *r* = ±1), or jointly otherwise. We propose to aggregate the resulting *p*-values *p*_*u*_ and *p*_*v*_ for 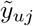 and 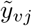 using Cauchy combination.[22] The Cauchy combination aggregates correlated *p*-values, and in this setting the test statistic can be written as:

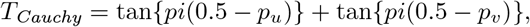

where *pi* is the mathematical constant approximately equal to 3.14159. The combined *p*-value for the tissue is approximated by:

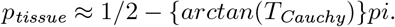

The *p*_*tissue*_ tests whether any association exists in the tissue. Unlike PrediXcan that tests associations averaged across all cell types, the *p*_*tissue*_ in MiXcan accumulates signals from different cell types. In transcriptome-wide studies, *p*_*tissue*_ from genes with cell-type-specific prediction models and *p*-values from genes with nonspecific models can be jointly used to adjust for multiple testing, and infer transcriptome-wide significant discoveries.

Note that *p*_*u*_ and *p*_*v*_ are building blocks of *T*_*Cauchy*_ and the resulting *p*_*tissue*_ is between *p*_*u*_ and *p*_*v*_. After the tissue-level hypothesis test, *p*_*u*_ and *p*_*v*_ can provide information on the cell type(s) driving the association. For example, *p*_*u*_ *<< p*_*v*_ indicates that a significant *p*_*tissue*_ is primarily driven by epithelial cells.

### Build MiXcan prediction models using GTEx mammary tissue data

MiXcan gene expression prediction models were developed using the GTEx v8 genotype and gene expression data for mammary tissue samples from 125 European ancestry women (dbGaP accession number phs000424.v8). PredictDB (http://predictdb.org) provides tissue-level expression prediction models trained on 337 men and women of European ancestry with mammary tissue data available in GTEx v8. PredictDB included mammary tissue elastic-net models for a total of 6461 genes that were well predicted by genetic variants (176,983 SNPs). An additional 1,715 SNPs were included in PredictDB mammary tissue MASHR models.[48] For the purpose of comparison to PrediXcan as a proof of concept, we developed cell-type-specific prediction models for these 6461 genes using all 178,698 SNPs in the PredictDB mammary tissue database. MiXcan cell-type-specific prediction models were developed for mammary epithelial cells, the cell of origin for breast carcinoma, and stromal (non-epithelial) cells.

#### *π* estimation

The epithelial cell proportion ***π***_*i*_ was estimated using 126 epithelial cell signature genes[18] available in the training dataset of 125 female mammary tissue samples. We first applied xCell[18] to estimate enrichment scores for multiple cell types, and then re-scaled the xCell epithelial cell enrichment score to range from 0 to 1 for use as a prior in MiXcan. The ***π***_*i*_ estimation was performed using 100 bootstrap samples (80% random draw with replacement). The final estimate was computed by excluding the most extreme 5% of bootstrap estimates in each of the two tails and averaging the remaining estimates.

#### Prediction model

Using 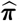, we modeled the cell-type-specific (or nonspecific) expression levels for each of the 6461 genes using MiXcan with tuning parameter λ selected by 10-fold CV. We adjusted for covariates that were used in GTEx eQTL analyses including age, platform, PCR, genomic principal components (PC) 1-5, and PEER factors 1-15.[49] For genes with non-zero 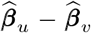, we performed ordinary least squares regression on the pre-selected variables for 200 bootstrap samples, and calculated the 95% bootstrap CI of 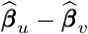. If the 95% CI excluded 0, we used cell-type-specific models with parameters estimated using the full data; otherwise, we used nonspecific models that were the same as PrediXcan.

### Evaluate MiXcan prediction accuracy in independent TCGA data

We evaluated the prediction performance of MiXcan in an independent dataset of 103 European ancestry female breast cancer patients with adjacent normal tissue samples from TCGA.[50, 51] To minimize the study effect, we re-processed the TCGA gene expression data using methods analogous to those used to process the GTEx expression data (https://gtexportal.org/home/documentationPage). Briefly, we required genes to have Transcripts Per Kilobase Million (TPM) > 0.1 in at least 20% of samples, and at least six reads in at least 20% of samples, resulting in a set of 25,702 out of 25,849 total genes that met these quality control (QC) requirements. Expression data were then normalized using the trimmed mean of M values method (TMM)[52] as implemented in the R package edgeR,[53] and the results were quantile-normalized to a standard normal distribution with mean=0 and variance=1. Comparison of these normalized gene expression levels showed no systematic differences between the GTEx and TCGA data (**Supplementary Fig. 7**). To process the genotype data, we removed all indels, monomorphisms, and ambiguous pairs (e.g. A/T, C/G). SNPs with > 5% missing genotypes or Hardy–Weinberg equilibrium (HWE) test *p*-value < 1e-05 were also removed. The remaining SNPs were aligned to build 37 coordinates, and imputation was performed on the TOPMed imputation server.[54] A total of 97% (54,663 out of 56,531) and 97% (52,031 out 53,876) SNPs used in MiXcan and PrediXcan prediction models were available for analysis.

We estimated the epithelial cell proportion in the TCGA samples as described above, and used this estimate to combine the predicted cell-type-specific gene expression levels from epithelial and stromal components into the tissue level. To evaluate predication accuracy, we computed the Pearson correlation between the predicted and observed tissue-level gene expression, and compared the results with the predicted tissue-level expression using PrediXcan. The observed bulk tissue expression levels showed significantly higher correlation with the tissue-level expression predicted by MiXcan compared with PrediXcan. To investigate the sources of the improved performance, we compared five approaches for predicting tissue-level gene expression:

- Existing PredictDB elastic-net models (PredictDB)
- PrediXcan trained on the same dataset as MiXcan (PrediXcan)
- Prediction model including interactions between SNPs and the xCell score (xCell interaction)
- Prediction model including interactions between SNPs and the MiXcan cell-type proportion (MiXcan_0_)
- Cell-type-specific prediction models with symmetric penalization (MiXcan).

These comparisons evaluated incorporating cell-type composition, use of the MiXcan cell-type proportion estimate, and symmetric penalization of the two cell types in prediction models, as well as use of a larger training set including both men and women in PredictDB. It is worth noting that “MiXcan_0_” and “xCell interaction” models are not applicable to GWAS datasets that lack cell-type composition information for the tissue of interest, and are included here only for the purpose of understanding the sources of improved prediction performance for MiXcan.

### Simulation studies

To evaluate the type I error and power of MiXcan association tests, we performed extensive simulation studies under a broad range of realistic models for the associations of genetic variants with gene expression (SNP-exp) and gene expression with disease (exp-disease). Mimicking real data, in each simulation, we generated a training dataset for building the expression prediction models, and a GWAS dataset for testing the associations of genetically predicted gene expression with disease.

For the training dataset, we simulated 300 bulk tissue samples with observed SNP genotypes and tissue-level gene expression. We assumed each tissue *i* ∈ (1, …, 300) was a mixture of two cell types and that the minor cell type (cell type 1) comprised an average of 40% of the tissue, with proportion *π*_*i*_ ∼ *Beta*(*α* = 2, *β* = 3). We further simulated the genotypes of 50 neighboring SNPs ***x***_*i*_ = {*x*_1*i*_, …, *x*_50*i*_} using the genome simulator R package *sim1000G*, with its default reference genome region (chromosome 4) and minor allele frequency (MAF) range 0.05–0.50. For the expression levels in the two cell types *u*_*i*_ and *v*_*i*_, we considered a linear additive model such that *u*_*i*_ = *β*_0_ + ***β***_1_***x***_*i*_ + *e*_*ui*_ and *v*_*i*_ = ***β***_2_***x***_*i*_ + *e*_*vi*_ where *e*_*ui*_, *e*_*vi*_ ∼ *N*(0, 1). The parameter *β*_0_ determines the mean expression difference in the two cell types under ***x***_*i*_ = 0, and ***β***_1_ and ***β***_2_ determine the association patterns between the SNPs *X* and gene expression level *Y* in the minor and major cell types, respectively. Then, the tissue-level gene expression is a weighted average of the expression levels in the two cell types: *y*_*i*_ = *π*_*i*_*u*_*i*_ + (1 − *π*_*i*_)*v*_*i*_.

We considered two SNP-exp settings: *Homogeneous SNP-exp Association* (***β***_1_ = ***β***_2_) and *Heterogeneous SNP-exp Association* (***β***_1_ ≠ ***β***_2_). Under the *Homogeneous SNP-exp* setting, we randomly selected one genetic variant *p* to be associated with expression levels in the two cell types and let *β*_1*p*_ = *β*_2*p*_ = 2 or -2 with equal chance. We altered *β*_0_ from -5 to 5 to evaluate the impact of the intercept (mean expression difference in two cell types under ***x***_*i*_ = 0) on TWAS. Under the *Heterogeneous SNP-exp* setting, we randomly selected two SNPs *p*_1_ ≠ *p*_2_ ∈ (1, …, 50) with SNPs *p*_1_ and *p*_2_ associated with expression levels in the minor and major cell types, respectively. Similar to the *Homogeneous SNP-exp* setting, we first evaluated the impact of the intercept by altering *β*_0_ from -5 to 5 while fixing 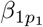 at 2 or -2 and 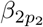 at 2 or -2 with equal chance. Second, we altered the magnitude of *β*_1*p*_ from 0 to 5 (allowing a random sign with equal chance), while fixing *β*_0_ at 2 and *β*_2*p*_ = ±2 to understand the impact of the SNP-exp association strength in the minor cell type. Third, we altered the magnitude of *β*_2*p*_ from 0 to 5 (allowing a random sign with equal chance), while fixing *β*_0_ at 2 and *β*_1*p*_ = ±2 to understand the impact of the SNP-exp association strength in the major cell type. Finally, to evaluate the impact of the sample size of the training dataset, we assessed sample sizes ranging from 100 to 300, while fixing *β*_0_ at 2 and the magnitude of non-zero components of ***β***_1_, ***β***_2_ at 2.

For the GWAS dataset, we simulated 500 samples with observed SNP genotypes and disease status for testing the associations of predicted gene expression levels with the disease. We assumed that the unobserved cell-type composition and cell-type-specific gene expression levels in this dataset followed the same distributions as in the training dataset. Disease risk was simulated using a logistic model, such that *logitP* (*d*_*j*_ = 1) = *η*_0_ + *η*_1_*u*_*j*_ + *η*_2_*v*_*j*_ for *j* ∈ (1, …, 500). The intercept *η*_0_ was set so that the disease prevalence was 50% in the samples. We considered five different settings for *η*_1_, *η*_2_ to capture the dynamic relationship between gene expression levels in the two cell types and disease risk:

- *No Exp-Disease Association*: *η*_1_ = *η*_2_ = 0, i.e. disease is not associated with the gene expression in either cell type.
- *Homogeneous Exp-Disease Association*: *η*_1_ = *η*_2_ = 0.2, i.e. disease is associated with the gene expression in both cell types in the same way.
- *Exp-Disease Association in Major Cell* : *η*_1_ = 0 and *η*_2_ = 0.2, i.e. disease is associated with the gene expression in the major cell type (cell type 2).
- *Exp-Disease Association in Minor Cell* : *η*_1_ = 0.2 and *η*_2_ = 0, i.e. disease is associated with the gene expression in the minor cell type (cell type 1).
- *Exp-Disease Association in Opposite Directions*: *η*_1_ = −0.2 and *η*_2_ = 0.2, i.e. disease is associated with the gene expression in the two cell types in opposite directions.

We compared the prediction accuracy, type I error, and power of MiXcan with PrediXcan, which ignores cell-type heterogeneity. The performance of MiXcan depends on estimates of the cell type proportion, 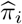. To evaluate the impact of misspecifying 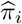, we let 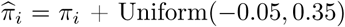 and rescaled it to (0,1), which resulted in a mean shift of the minor cell type proportion from 40% to 45% and reduced correlation with the true *π*_*i*_ to 0.9. We compared the results of MiXcan, using either the true or misspecified cell type proportion, with PrediXcan in 200 Monte Carlo simulations.

### Apply MiXcan to perform cell-type-specific TWAS of breast cancer

#### Cell-type-specific TWAS of breast cancer

We performed TWAS of breast cancer risk using GWAS data from the Discovery, Biology, and Risk of Inherited Variants in Breast Cancer (DRIVE) study. Genotype data for 60,014 women (32,438 cases and 27,576 controls) assayed on the Oncoarray[24] were downloaded from dbGaP (phs001265.v1.p1), which provides excellent coverage of most common variants. After imputation (as described above for TCGA data), 95% (53,528 out of 56,531) and 95% (51,049 out of 53,876) SNPs used in MiXcan and PrediXcan prediction models were available for analysis, respectively.

Principle component analysis (PCA) was performed using 20,629 SNPs, after excluding SNPs with a missing rate above 0.01% and selecting SNPs in approximate linkage equilibrium using PLINK[55] (indep-pairwise option with window size=50kb, step size=5, *r*^2^ threshold=0.05). EIGENSOFT v6.1.4 was used to compute PCs with the fast mode option enabled, which implements the FastPCA approximation.[56] The first PC separated individuals of African (e.g. from Nigeria, Uganda and Cameroon) vs. European (e.g. from Australia) ancestry. In total, 58,648 women (31,716 cases and 26,932 controls) of European ancestry determined by PCs were included in TWAS analyses.

MiXcan, PrediXcan and PredictDB elastic-net mammary tissue models were applied to the individual-level genotype data to perform cell-type-specific or tissue-level TWAS, as described above. All three models were adjusted for the same covariates, including age, country of origin and the top 10 PCs.[15]

#### Evaluation of TWAS findings

We evaluated significant TWAS genes identified by MiXcan and PrediXcan in a substantially larger study of 228,951 European ancestry women (122,977 cases and 105,974 controls) from the combined DRIVE and Breast Cancer Association Consortium (BCAC) GWAS meta-analysis of breast cancer.[11] The summary statistics for the “Combined Oncoarray, iCOGS GWAS meta analysis” were downloaded from https://bc ac.ccge.medschl.cam.ac.uk/bcacdata/oncoarray/oncoarray-and-combined-summary-result/gwas-summary-results-breast-cancer-risk-2017/. Associations of predicted tissue-level gene expression with breast cancer risk were evaluated using S-PrediXcan[27] with PredictDB[23] elastic-net models derived from GTEx v8 mammary tissue data for 337 men and women of European ancestry.

We also determined whether TWAS genes identified by MiXcan and PrediXcan were located within 500 kb of 214 previously reported genome-wide significant breast cancer susceptibility loci.[11, 12, 13]

### MiXcan software

We developed a computationally efficient R package *MiXcan* to facilitate estimation of cell-type-specific prediction models for genetically regulated gene expression in two cell components of bulk tissue data, and to perform cell-type-specific TWAS. The *MiXcan* R package, and pre-trained models for the epithelial and stromal (non-epithelial) cell components of mammary tissue derived from 125 European ancestry women in GTEx v8 are freely available at https://github.com/songxiaoyu/MiXcan.

## Acknowledgements

This study was supported by grants from the National Institutes of Health (R01CA237541, U24CA210993, P30CA196521, R01CA264987, R01CA166827, R01CA168893) and by the computational resources and staff expertise provided by Scientific Computing at the Icahn School of Medicine at Mount Sinai. The BCAC breast cancer genome-wide association analyses were supported by the Government of Canada through Genome Canada and the Canadian Institutes of Health Research, the ‘Ministère de l’Économie, de la Science et de l’Innovation du Québec’ through Genome Québec and grant PSR-SIIRI-701, the National Institutes of Health (U19 CA148065, X01HG007492), Cancer Research UK (C1287/A10118, C1287/A16563, C1287/A10710) and the European Union (HEALTH-F2-2009-223175 and H2020 633784 and 634935); all studies and funders are listed.[11]

## Competing interests

E.J. is an employee at Regeneron. The remaining authors declare no competing interests.

## Supplementary Figures

**Supplementary Figure 1.**
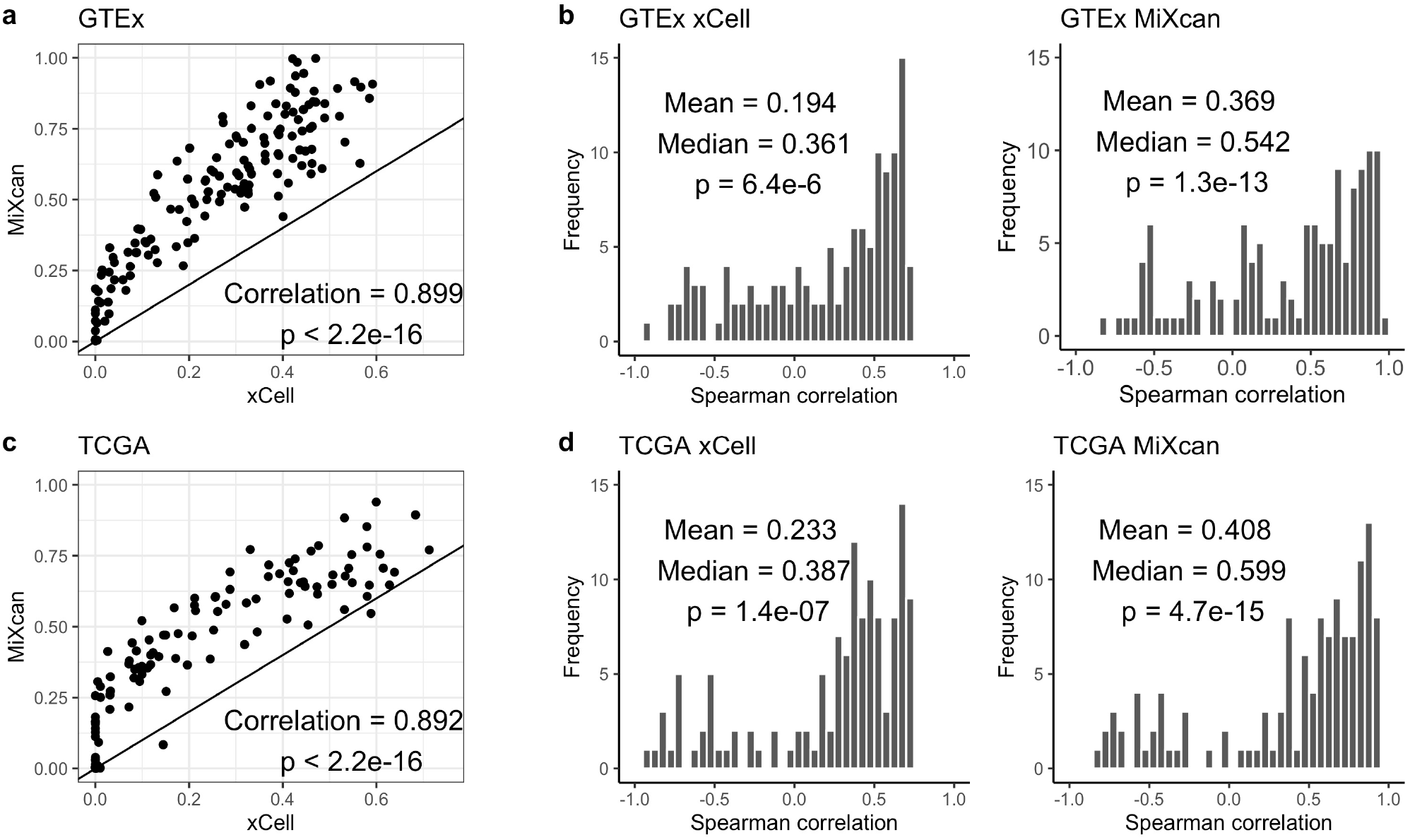
Epithelial cell proportion estimates. The MiXcan epithelial cell proportion and xCell epithelial cell enrichment score were estimated in normal mammary tissue samples from European ancestry women in GTEx (N=125) and TCGA (N=103). The estimated xCell score and MiXcan proportion were highly correlated in both the GTEx (**a**) and TCGA (**c**) samples. Overall, the MiXcan proportion was more highly correlated than the xCell score with the measured expression levels of 126 genes in the xCell epithelial cell gene signature, and the correlations were significant in both the GTEx (**b**) and TCGA (**d**) samples.

**Supplementary Figure 2.**
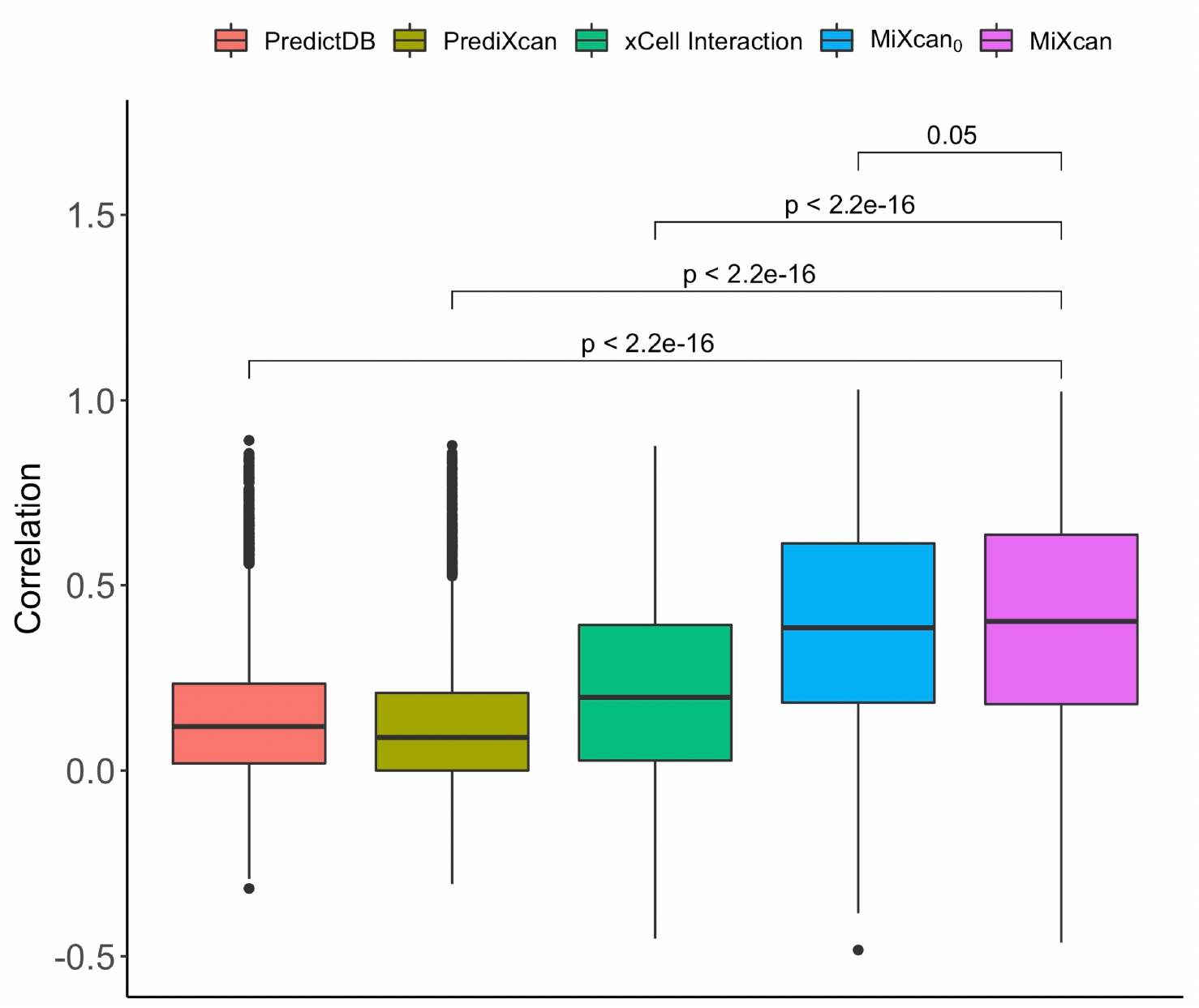
Gene expression prediction accuracy in independent validation dataset. Correlation of true and predicted tissue-level gene expression was evaluated for 6461 genes in an independent dataset of adjacent normal mammary tissue samples from 103 European ancestry women with breast cancer in TCGA. The PredictDB (median *r*=0.12) database includes the tissue-level gene expression prediction weights for elastic net models trained using mammary tissue samples from 337 European ancestry men and women in GTEx v8. PrediXcan (median *r*=0.10) uses the same approach as PredictDB but is trained only on the subset of 125 European ancestry women used to train the other prediction methods. xCell Interaction (median *r*=0.20) is an elastic net model that includes interactions between genetic variants and the xCell epithelial cell enrichment score. MiXcan_0_ (median *r*=0.38) is an elastic net model that includes interactions between genetic variants and the estimated epithelial cell proportion. MiXcan (median *r*=0.41) is the final symmetric penalization approach for estimating epithelial and stromal cell models that can be applied to GWAS datasets without cell-type proportion information.

**Supplementary Figure 3.**
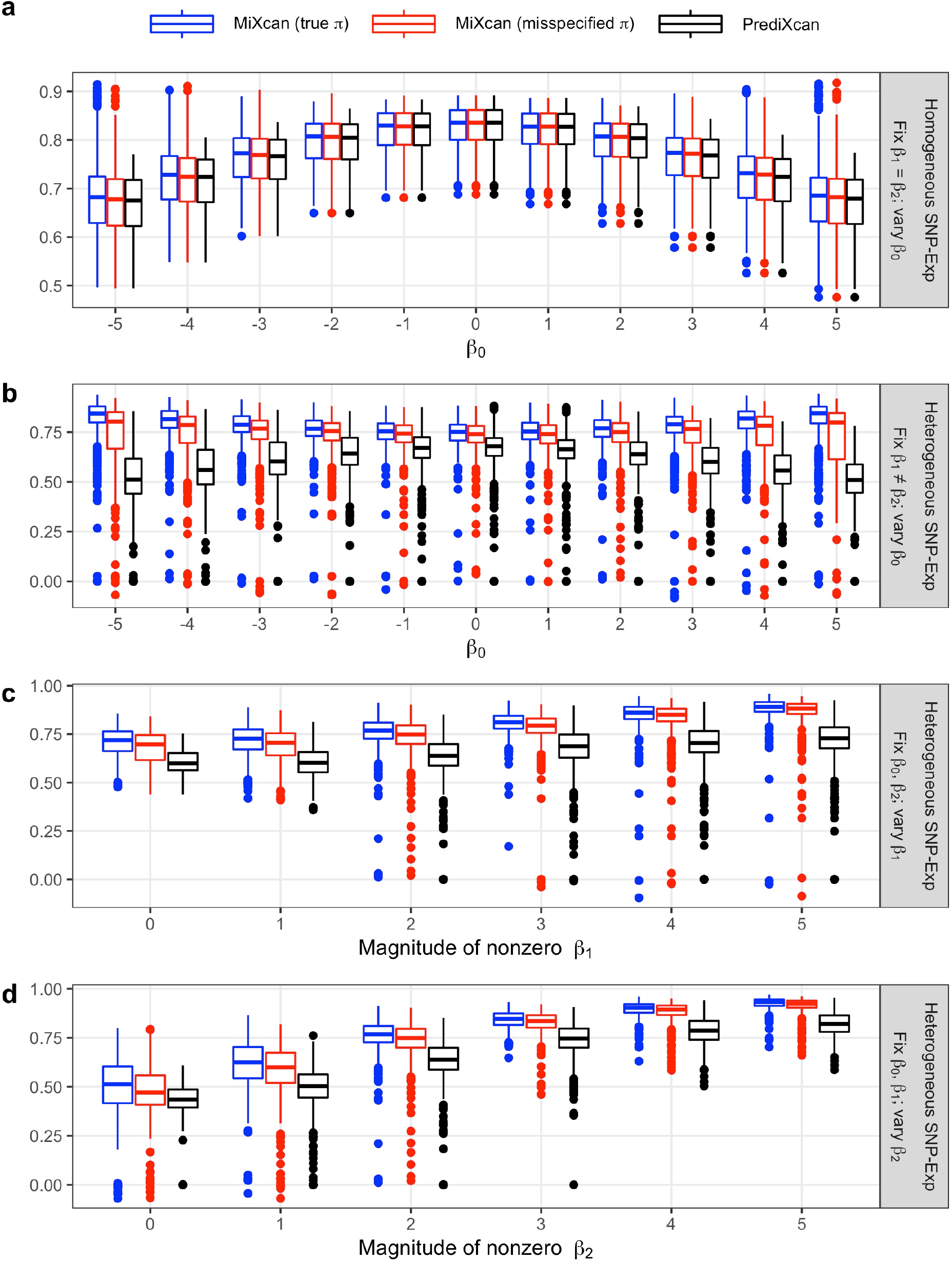
Simulation studies to evaluate gene expression prediction accuracy. Correlation of true and predicted tissue-level gene expression using the MiXcan or PrediXcan approaches in independent test sets (N=500) simulated under a range of realistic data scenarios as shown in Figure 3.

**Supplementary Figure 4.**
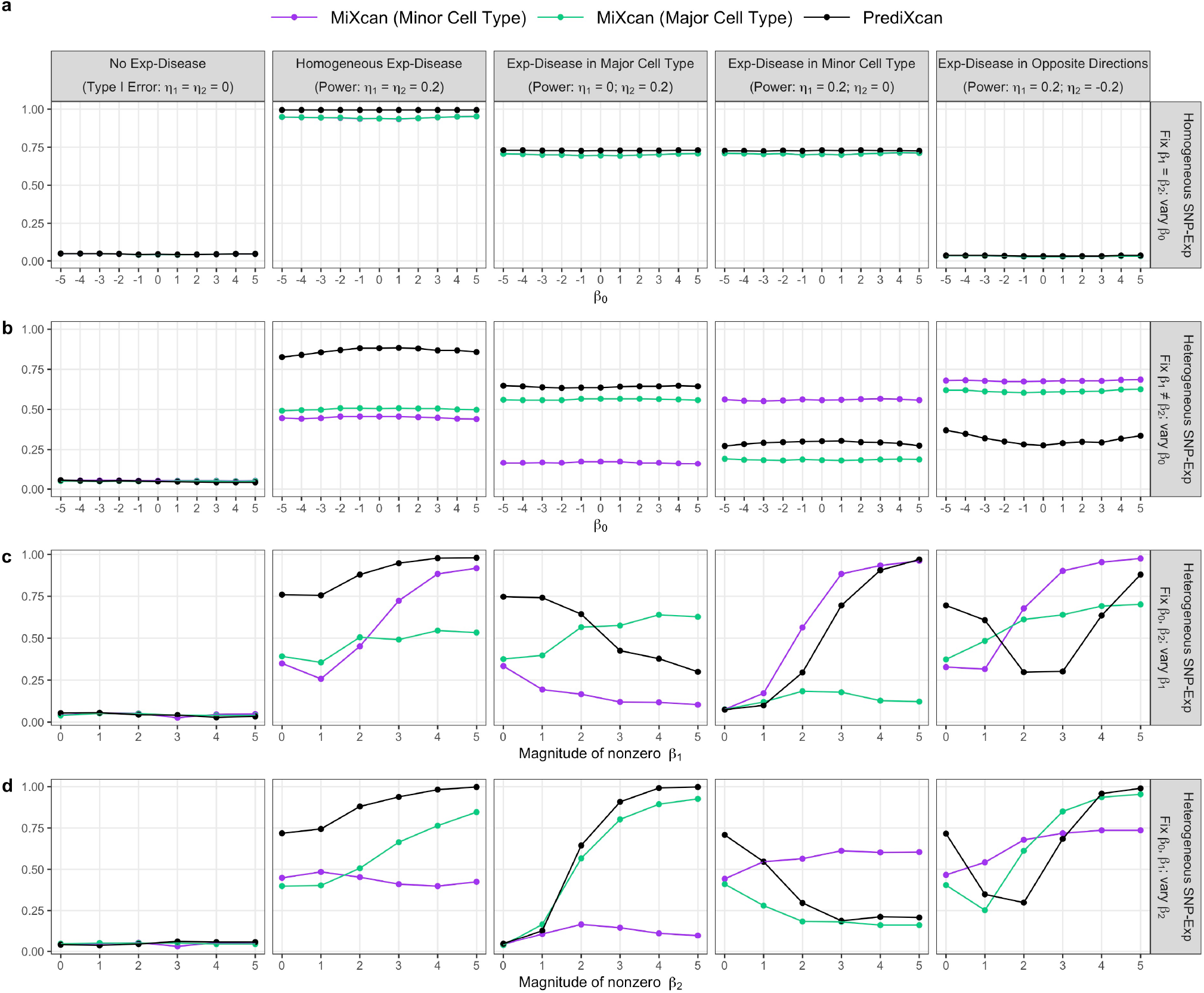
Simulation studies to assess cell-type-specific associations. Type I error and power of MiXcan to detect cell-type-specific associations of gene expression with the disease in the minor and major cell types when the cell-type proportion is accurately estimated. Independent test sets (N=500) were simulated under a range of realistic data scenarios as shown in Figure 3.

**Supplementary Figure 5.**
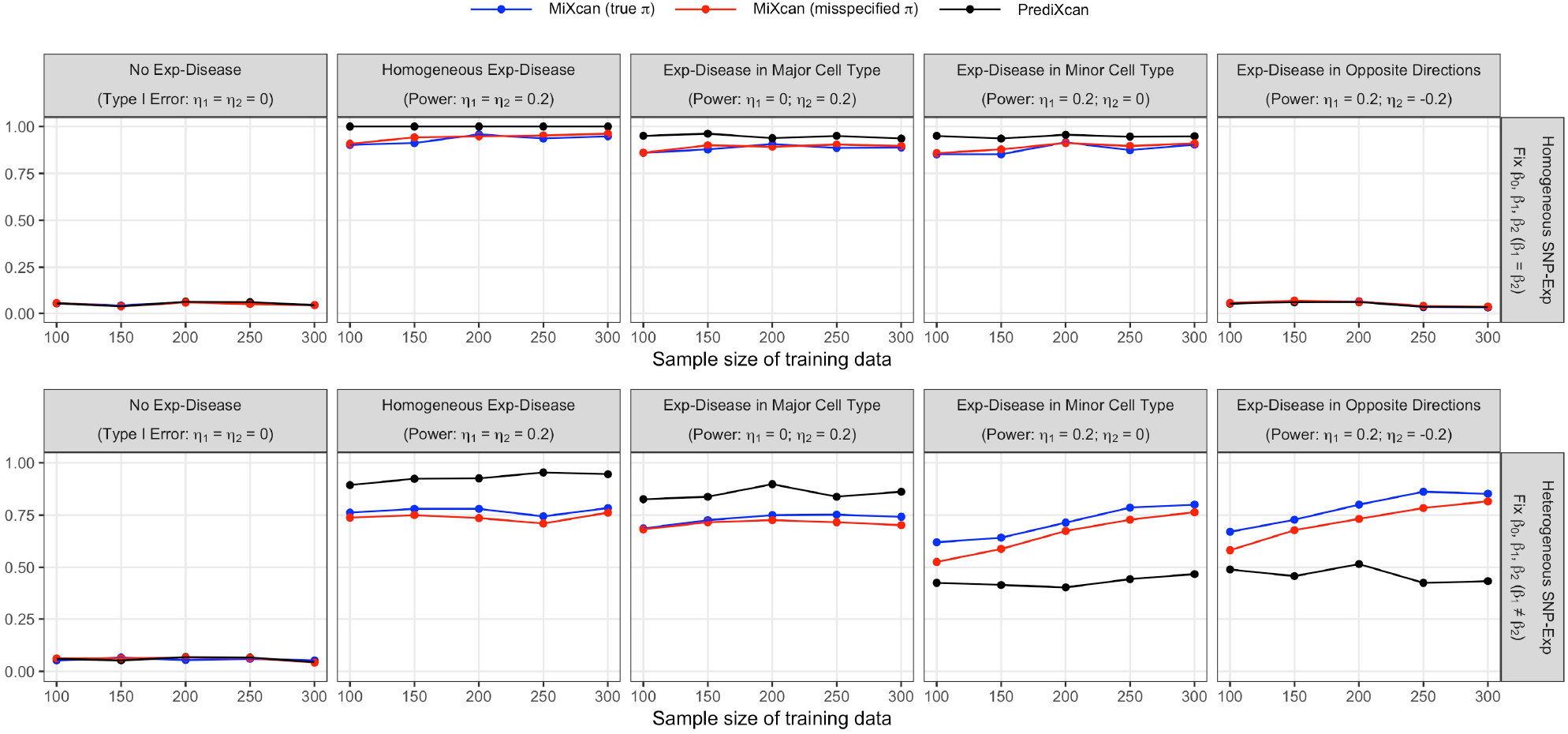
Simulation studies to assess the impact of the training data sample size. Type I error and power of the MiXcan or PrediXcan approaches trained using 100-300 samples were evaluated in independent test sets (N=500) simulated under a range of realistic data scenarios as shown in Figure 3.

**Supplementary Figure 6.**
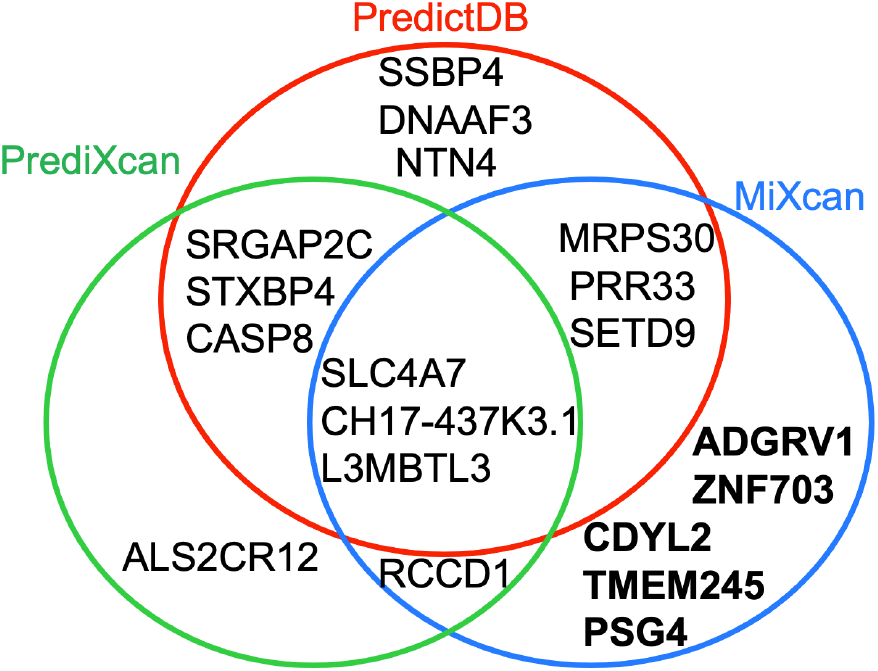
Transcriptome-wide association studies. Genes significantly associated with breast cancer using PredictDB, PrediXcan, and MiXcan at *P <* 7.7 × 10^−6^ applying a Bonferroni correction for the 6461 genes tested in 31,716 breast cancer cases and 26,932 controls of European ancestry from the DRIVE study (dbGaP phs001265.v1.p1). The PredictDB elastic net models were trained using mammary tissue samples from 337 men and women, whereas PrediXcan and MiXcan were trained using only 125 women of European ancestry in GTEx v8.

**Supplementary Figure 7.**
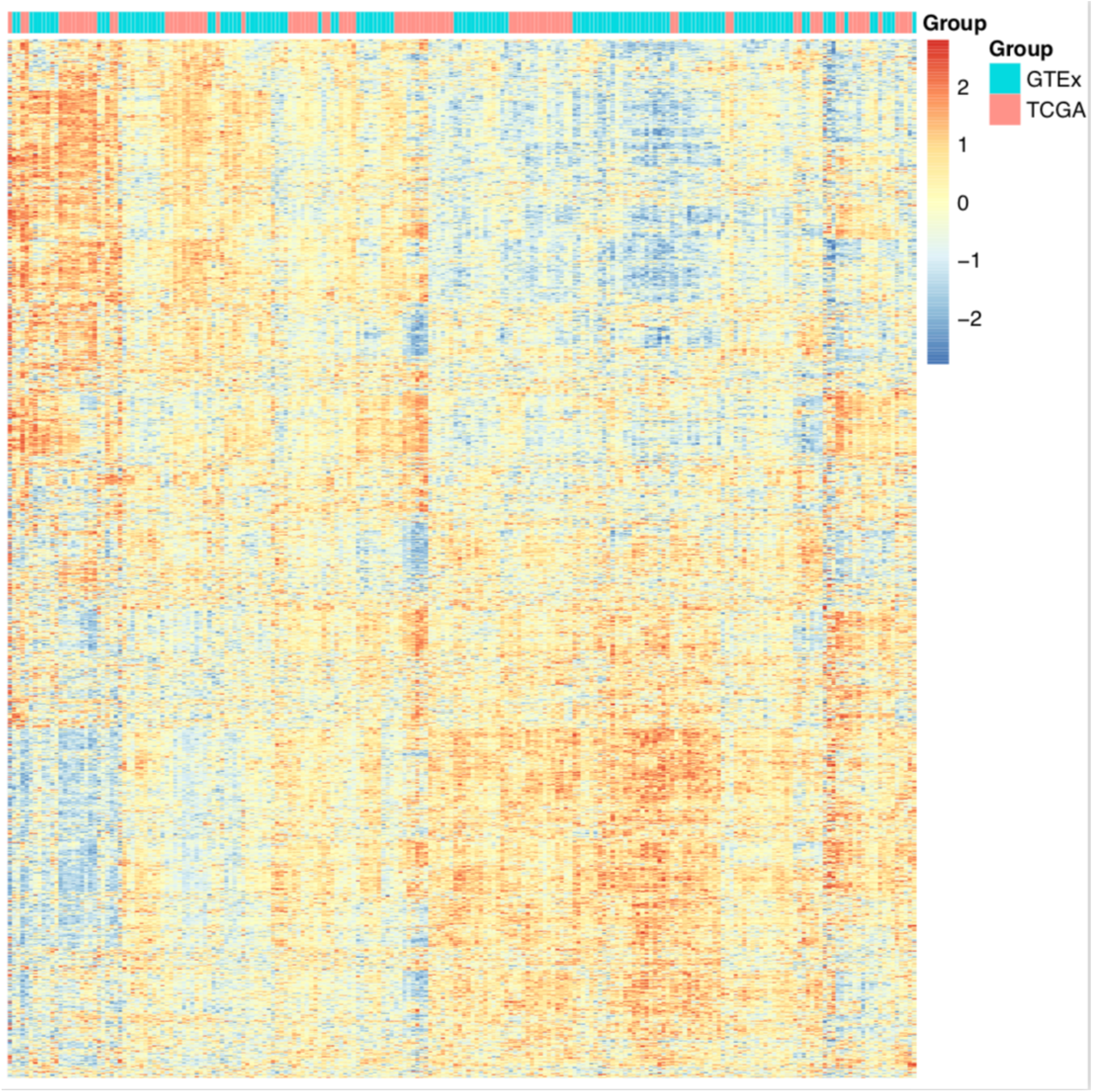
Heatmap of mammary tissue gene expression levels. There were no systematic differences between the normalized gene expression levels for 6421 genes present in both the GTEx (N=125) and TCGA (N=103) normal mammary tissue datasets, which were used for training and independent validation, respectively.

